# OPTRACE: Optical Imaging–Guided Transplantation and Tracking of Cells in the Mouse Brain

**DOI:** 10.1101/2025.05.24.655973

**Authors:** Jinghui Wang, Colleen Russel, Mikolaj Walczak, Dawei Gao, Guanda Qiao, Xiaoxuan Fan, Chengyan Chu, Miroslaw Janowski, Piotr Walczak, Yajie Liang

## Abstract

Intracerebral cell transplantation holds great promise for treating stroke and other neurological disorders, but progress is hindered by challenges in precise cell delivery and limited methods for monitoring post-engraftment dynamics. To address these limitations, we introduce OPTRACE (OPtical imaging-guided Transplantation and tRAcking of CElls), a two-step framework that integrates precision delivery with longitudinal in vivo tracking to advance cell therapy. During transplantation, OPTRACE enables real-time visualization using cost-effective translucent glass micropipettes, guided by predictive modeling to optimize delivery parameters. A novel pulse-elevation injection technique further enhances the precision of superficial cortical engraftment. Following transplantation, multicolor labeling combined with two-photon fluorescence microscopy permits longitudinal single-cell tracking, revealing host microglial responses, and altered neuronal calcium signaling at the graft interface. By unifying high-resolution delivery and tracking strategies, OPTRACE provides a powerful platform for elucidating engraftment biology and accelerating the development of effective cell-based interventions for neurological diseases.

## Introduction

Cell therapies are being actively explored, both preclinically and clinically, for the treatment of stroke, brain injury, and various neurodegenerative disorders ^1, 2, 3, 4, 5, 6^. However, the therapeutic potential of these cells depends critically on their precise and effective delivery^7, 8^. Intracerebral transplantation typically involves injecting high-density cell suspensions into damaged brain regions using a syringe-attached needle. Phase I clinical trials have demonstrated the procedural safety of intracerebral cell implantation, with no significant adverse effects reported ^9, 10, 11^. Despite this, cell retention and viability remain low, with studies reporting survival rates of as little as 3% ^7, 12, 13, 14^. While post-transplantation survival can be influenced by factors such as inflammation, lack of trophic support, or an unfavorable extracellular matrix, cell damage during injection and leakage from the injection site may also contribute to poor outcomes ^7, 12, 14^. Notably, there is limited guidance on predicting cell retention or growth outcomes based on injection depth or implantation volume during intracerebral cell delivery.

To improve cell retention during delivery, a critical step in successful cell transplantation is essential to closely monitor the injection process for optimizing delivery parameters accordingly. A major barrier to clinical translation has been the lack of noninvasive methods to assess whether appropriate cell doses are being administered ^15, 16^. Even in preclinical studies, reliable approaches for monitoring the cell delivery process remain limited ^17^. On the other hand, in the post-transplantation phase, there remains a critical need for noninvasive imaging techniques capable of tracking the survival, engraftment, migration, and distribution of transplanted stem cells in vivo with high spatial and temporal resolution ^18, 19^. Most existing studies on stem cell therapy rely on histological analysis at a few discrete timepoints to assess outcomes ^20, 21, 22^, which fails to capture the dynamics of implanted cells, or interactions between graft and host cells. A central challenge in advancing cell transplantation therapy is the limited understanding of donor cell fate in living animals—specifically their morphology, function, and interactions with host neurons and glial cells over time. Without appropriate tools, this post-transplantation phase remains a “black box” in need of illumination.

In this study, we present OPTRACE (OPtical imaging-guided Transplantation and tRAcking of CElls), a two-step framework leveraging optical imaging to overcome critical challenges in cell transplantation and post-transplantation phases. During transplantation, OPTRACE utilizes transparent glass micropipettes as injectors for real-time visualization of the injection process, enabling precise optimization of injection depth and volume, validated by theoretical modeling. Notably, OPTRACE introduces the pulse-elevation technique that significantly improves cell retention in the superficial layers of the mouse cerebral cortex compared to conventional continuous injection methods. Post-transplantation, OPTRACE employs intravital two-photon fluorescence microscopy (TPFM) for high-resolution, longitudinal tracking of grafted cells, and dynamic monitoring of host neuronal and microglial responses at the graft site. Collectively, these integrated capabilities establish OPTRACE as a robust platform for addressing key barriers in cell transplantation and advancing regenerative cell therapies.

## Results

### Illuminating cell therapy through OPTRACE

OPTRACE integrates three core components to advance intracerebral cell transplantation (**Fig. 1**). First, cost-effective transparent glass pipettes enable direct visualization of cells during loading and injection, reducing procedural uncertainty. To enhance accessibility, we developed Affordable Pipette Pullers (APPs), a low-cost method for fabricating functional injectors, eliminating reliance on expensive specialized equipment. Second, cells are genetically labeled with fluorescent proteins (FPs) or bioluminescent reporters (e.g., luciferase) for precise quantification or tracking. Multiplexed FPs assign unique spectral signatures for high-fidelity single-cell tracking, as shown in our prior work ^23^. These two components are prepared at the pre-transplantation phase. The third component involves three optical imaging modalities: wide-field microscopy, bioluminescence imaging, and TPFM. For the transplantation phase, wide-field microscopy facilitates real-time cell visualization, while bioluminescence imaging quantifies cell retention and monitors growth to validate theoretical models in vitro. As a powerful tool increasingly used for high-resolution imaging of brain tissue^24, 25, 26^, TPFM supports both transplantation procedure (real-time confirmation of optimized delivery) and post-transplantation phases (longitudinal tracking of implanted cells in vivo), owing to its deep tissue penetration and subcellular resolution. The following sections detail OPTRACE’s implementation and its potential to revolutionize intracerebral cell transplantation.

**Figure 1.**
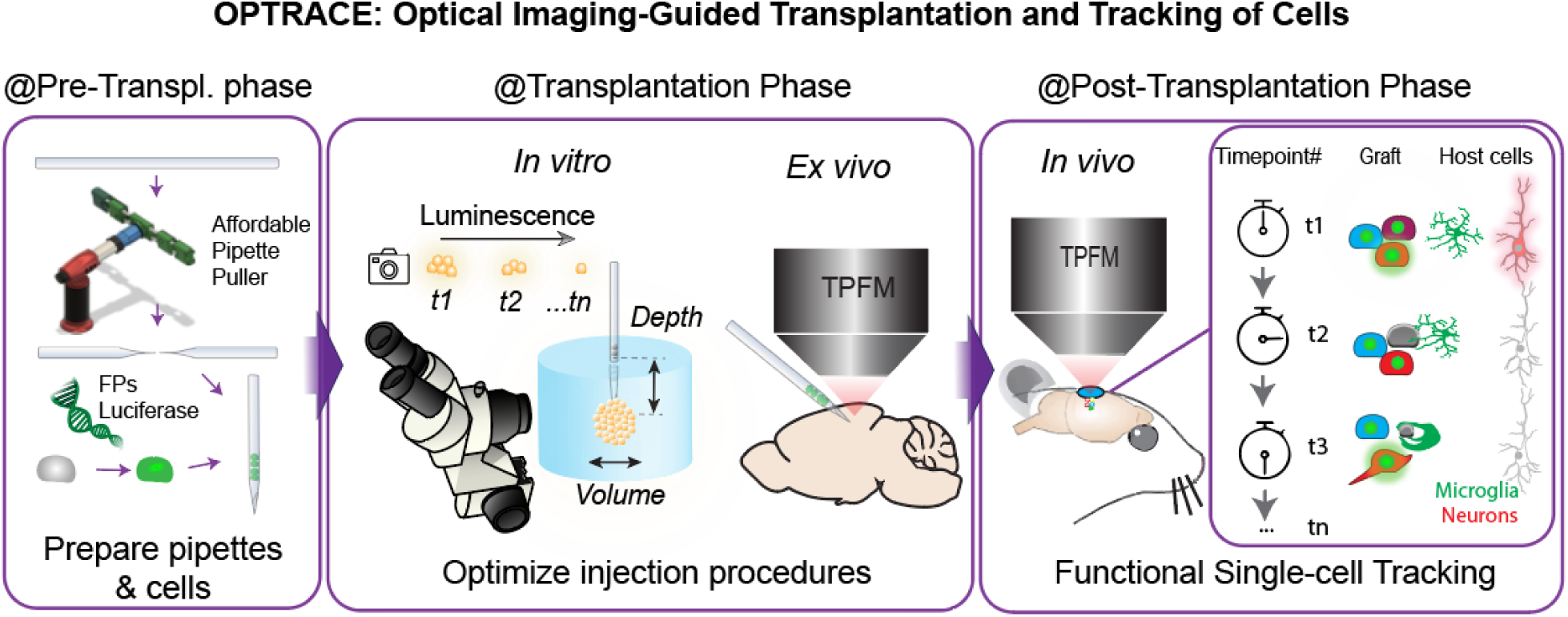
Schematic illustrating the application of OPTRACE (Optical Imaging-Guided Transplantation and Tracking of Cells). During the pre-transplantation phase, transparent glass pipettes are prepared for direct visualization of cells during loading and injection, and cells are genetically labeled with fluorescent reporters for quantification and tracking. In the transplantation phase, optical imaging enables real-time monitoring of the procedure and assessment of cell retention and viability. Post-transplantation, TPFM supports longitudinal tracking of implanted cells and reveals dynamic responses of host cells at the transplant site.

### Affordable Transparent Pipette Pulling and Real-Time Visualization Platform for Cell Transplantation in OPTRACE

To achieve OPTRACE, we utilized transparent glass pipettes as a replacement of metal needles, which forbid visualization of cells during cell injection. To overcome the cost barrier that limits access to commercial pipette pullers, we upgraded previous Tinkcad designs of APP ^27^ by revising them through Fusion 360 (Autodesk, **Fig. 2A**), enhancing both the robustness and consistency of the APP. The redesigned APP 2.0 features improved ergonomics, reliability, and ease of use (**Fig. 2B, upper panel**). Additionally, we developed a multi-channel APP 2.0 capable of simultaneously producing four pipettes for high-throughput fabrication (**Fig. 2B, lower panel,** design files are available here: https://github.com/liangy10/APP2). To improve usability, we also engineered a new torch adapter (**Fig. 2)** that securely attaches to the APP body via built-in threads, significantly improving handling and precision during pipette pulling. Its effectiveness is demonstrated in imaging results (**Fig. 2D**), where cells are visible within the glass pipette under both brightfield or fluorescence microscopy (indicated by arrows). More importantly, under both but not simultaneous wide-field microscopy and TPFM (**Fig. 2E**), we readily located pipettes (**Fig. 2F**, left panel) containing fluorescently labeled cells and monitored the injection process in real time under TPFM (**Fig. 2F**, right panel).

**Figure. 2.**
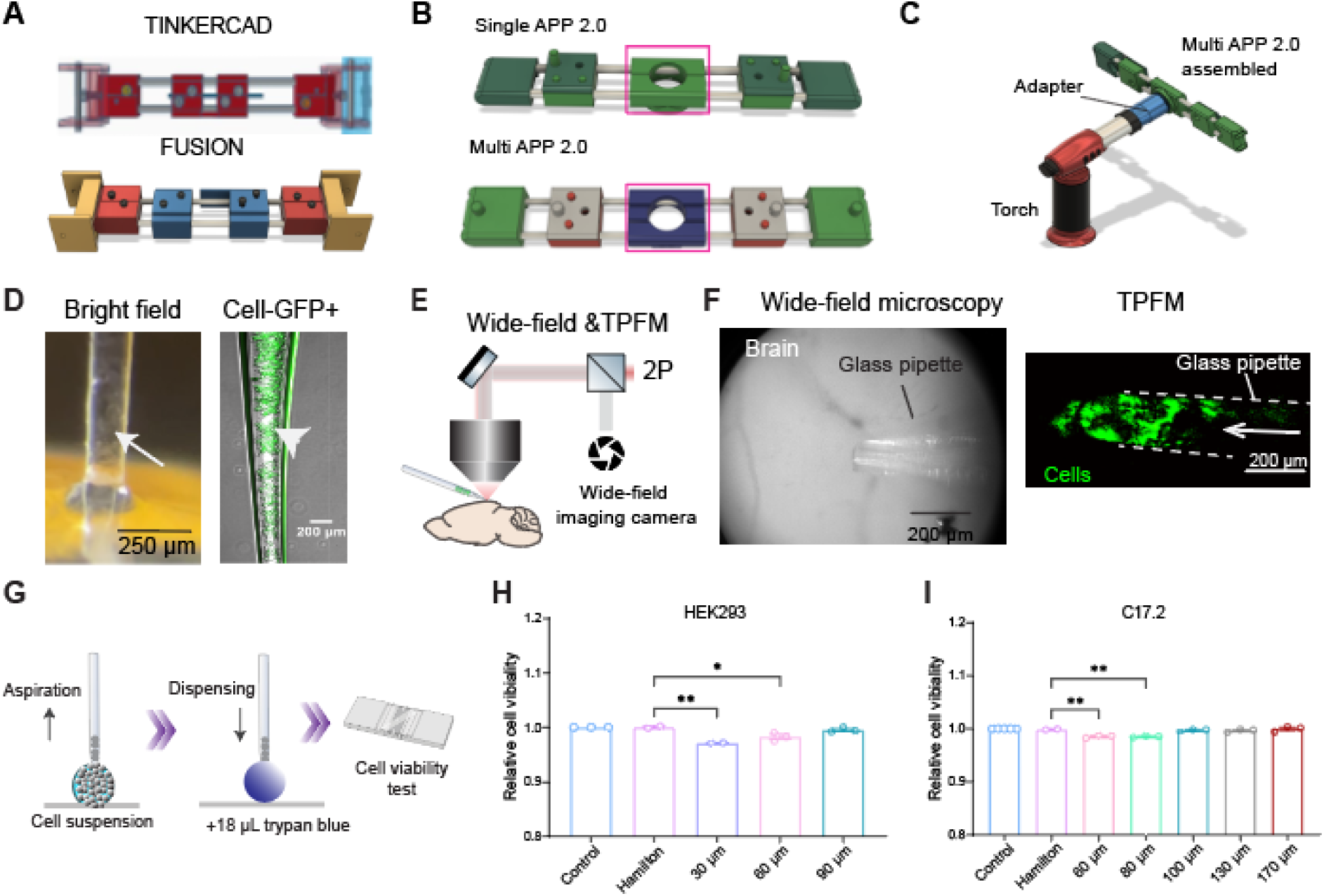
Upgraded APP for the production of the injector and determination of optimal injector diameter for cells. (**A**) APP designs in Tinkercad and in Fusion 360. (**B**) Upgraded APP 2.0 includes single-channel and multi-channel designs. (**C**) Assembled APP for use of pipette pulling. (**D**) Bright field and fluorescence channels imaging of glass pipette filled with cells labeled with green fluorescent protein (GFP). (**E**) Schematic of the ex vivo imaging setup. (**F**) Wide-field imaging (Left) and TPFM (Right) of a glass pipette filled with C17.2 cells labeled with GFP during injection into the mouse brain (ex vivo). (**G**). Workflow for testing the effect of flowing through the needle on the viability of cells. (**H, I**) Viability of HEK293 cells (**H**) or C17.2 (**I**) at different diameters of injectors. For (**H**): Hamilton vs 30 μm: p=0.0014; Hamilton vs 60 μm: p=0.0179, n=3,2,2,3,3. For (**I**): Hamilton vs 60 μm: p=0.0126; Hamilton vs 80 μm: p=0.0362; n=7,3,3,3,3,3,3. Differences were analyzed by one-way ANOVA followed by Tukey’s test. (\**p* < 0.05, *\*\* p* < 0.01).

Needle diameter plays a critical role in the success of delicate cell transplantation procedures ^8, 28, 29^. While small-diameter needles are favored for minimizing tissue damage, they can compromise cell viability due to mechanical stress during injection. Thus, we examined the relationship between needle diameter and cell viability after flowing through the needle. Target cells (HEK293 as a control or mouse neural stem cell line C17.2) were aspirated and subsequently dispensed through the glass pipette needles (**Fig. 2G**). Viability was evaluated, revealing significant reduction in the 30 µm (p = 0.0014) and 60 µm (p = 0.0179) groups compared to the Hamilton group for HEK293 cells (**Fig. 2H**), whereas the 90 µm pipette did not show a significant reduction (p = 0.7886, **Fig. 2H**). For C17.2 cells, the 60 µm (p = 0.0126) and 80 µm pipettes (p = 0.0362) resulted in significantly reduced viability (**Fig. 2I**), while no significant difference was observed for pipettes larger than 100 µm when compared to the Hamilton control (p = 0.328, Fig. 2I). Therefore, diameters in the range of 100–120 µm were used for subsequent experiments.

### Modeling Cell Retention Rate as a Function of Injection Depth for Optimizing Transplantation into the Brain

Research indicates that the survival rate of transplanted neural progenitor cells can be as low as 1-2%, with substantial cell loss post-transplantation ^30, 31, 32^. This highlights the importance of optimizing transplantation parameters. We modeled the relationship between the depth of injection (*d*) and the cell retention rate (*R,* **Fig. 3A**) with other fixed parameters, such as cell density, needle diameter, injection velocity, etc. Due to the viscosity of the brain tissue, there is a sealing vector (*N_s_*) pointing from the host tissue to the center of needle during the transplantation procedure. As more depth cells are injected inside the tissue (as *d* increases), there is more resistance against the cells preventing them from leaking out through the needle track. Therefore, the magnitude of this vector is in positive relation to the injection depth. For simplicity, assume it’s a linear or power relationship:*N*(*d*) = *k* ⋅ *d*^*n*^, where k>0, n≥1, in which n is adjustable to represent nonlinear growth. As retention rate *R* is positively related to sealing force, approaching 100%, we modeled it using a sigmoid or saturation function:*R*(*d*) = 100% • (1 − *e*^−*αN*^(*d*)). Substituting N and assuming n = 1, we got *R*(*d*) = 100% • (1 − *e*^−*αkd*^), where α is a scaling parameter related to how strongly sealing affects retention and *k* is the scaling coefficient for sealing force. If we merge the α and k into one constant *β*, we get R(d) = 100% • (1 − e^−βd^) or if in fraction,*R*(*d*) = 1 − *e*^−*βd*^ (Equation 1, **Fig. 3B**). This model intends to capture the increase in retention with depth and reflect how sealing forces work in soft, viscoelastic tissue. At depth *d* = 0, *R(d)* = 0, meaning no retention as cells are all leaking out. As depth increases, retention approaches 100% asymptotically. The parameter *β* controls how steeply retention increases with depth. A larger *β* means we get close to 100% retention with shallower injections (**Fig. 3B**), while a smaller *β* means we need to inject deeper to achieve high retention. As this is a simplified function, we use *β* as an integrated variable that is affected by tissue viscosity, expandability, engraft rigidity, and viscosity, needle shape, etc. But once transplantation settings are determined, such as target tissue, engraft, etc, *β* could be determined through experiment, which was entailed below.

**Figure. 3.**
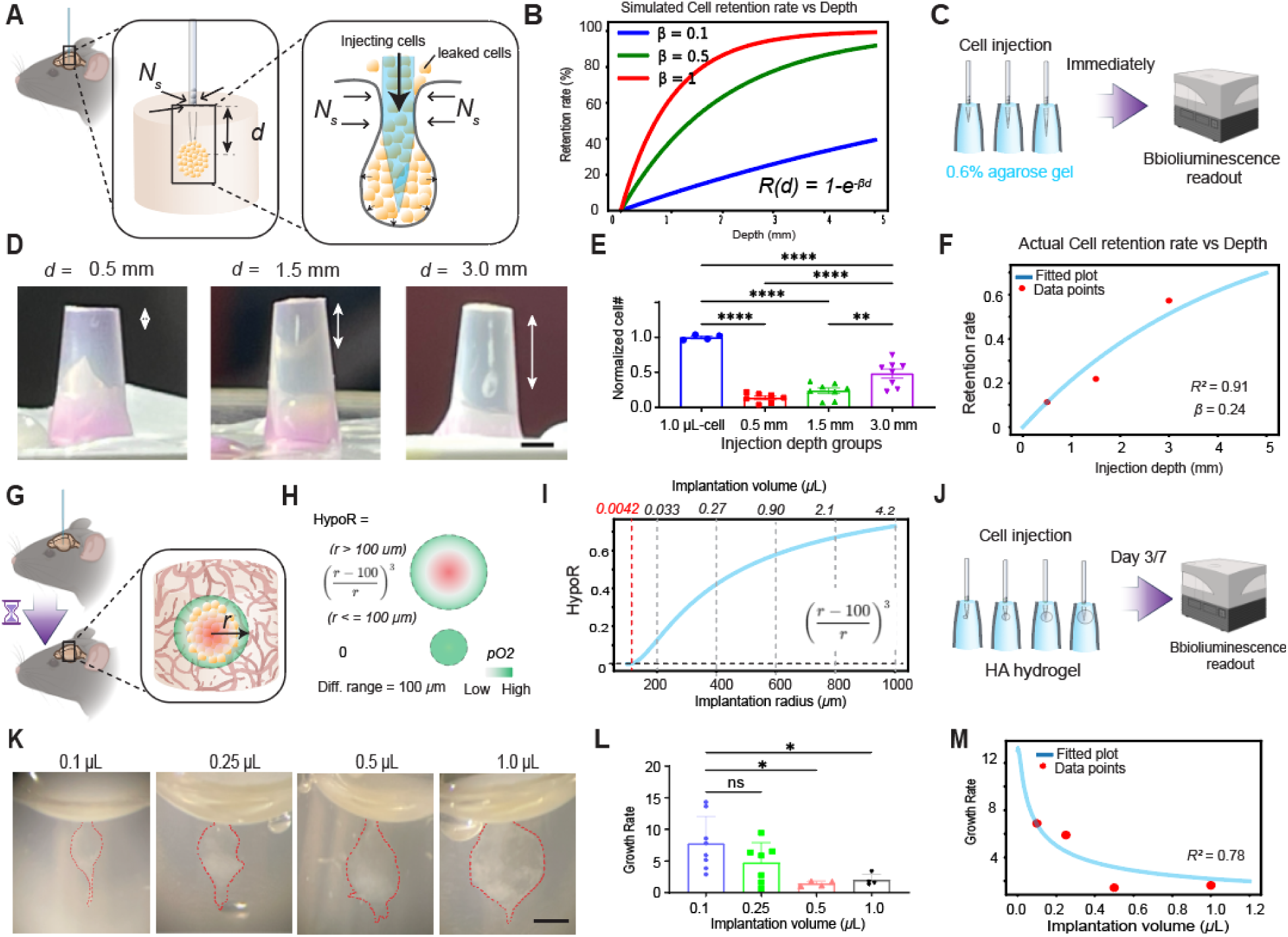
Modeling cell retention rate and hypoperfused cell fractions. (**A**) Schematic representation of a needle inserted into the brain, where sealing force *N_s_* is proportional to needle insertion depth *d*. (**B**) Simulated cell retention rate at different beta value. Experimental setup (**C**) for validating the model after injecting cells at different depths within the 0.6% low-melting agarose (**D**), using bioluminescence imaging to quantify retained cells (E, p < 0.0001, n = 4,7,8,8). (**F**) Fitting the theoretical curve with experimental data. (**G**) Schematic illustration of implanted cell mass undergoing hypoperfusion at post-transplantation phase. (**H**) Relationship between hypoperfused ratio and diameter of the implanted cell mass. (**I**) HypoR as a function of radius of implanted cell mass. Upper horizontal axis shows corresponding volume of implanted cell mass. (**J**) Experimental setup to verify the modeling. (**K**) Cells of different volumes were injected into hydrogel. (**L**) Quantification of growth rate after 4 days of culture. (0.5 µL, p = 0.0188), and 1.0 µL, p = 0.0343, n=8,7,4,4). (**M**) Fitting the theoretical curve with experimental data. Differences were analyzed by one-way ANOVA followed by Tukey’s test. (* p < 0.05, **p < 0.01, ****p < 0.0001)

To obtain experimental support for our modeling, we injected mouse neural stem cells (C17.2) into different depths of the 0.6% low-melting agarose gel (simulation of the brain tissue ^33, 34^) and quantified the fraction of cells that are retained through bioluminescence imaging (BLI) as C17.2 cells express luciferase (**Fig. 3C**). After injection, the needle tracks and cells were clearly visible (**Fig. 3D**). Quantification of cell number in each group revealed that, compared to the 1.0 μL cell suspension group, signals from the 0.5 mm, 1.5 mm, and 3.0 mm depth groups were significantly reduced (**Fig. 3E,** p < 0.0001). Further analysis revealed that the cell signal in the 3.0 mm group was significantly higher than in the 0.5 mm and 1.5 mm groups. However, there was no significant difference between the 0.5 mm and 1.5 mm groups (p = 0.4059, **Fig. 3E**). We then used this data to calculate the *β* value in the above Retention function and got the *β* value of 0.24 with good fitting (Pear’s square = 0.91, **Fig. 3F**). Thus, under our experimental conditions, the equation *R*(*d*) = 1 − *e*^−0.24*d*^ (Equation 2), serves as the retention rate prediction function. These findings confirm our modeling hypothesis and provide theoretical guidance for predicting cell retention rate as a function of injection depth.

### Modeling Hypoperfused Cell Fractions as a Function of Graft Volume

After cells are implanted into the brain, they experience stress due to limited oxygen and nutrient delivery (**Fig. 3G**), referred to as hypoperfusion ^2, 35^. Conceptually, the larger the implanted cell mass, the greater the fraction of cells that become hypoperfused. Under normal physiological conditions, the partial pressure of oxygen (pO₂) in the cerebral cortex ranges from 30 to 40 mmHg. When pO₂ drops below 5 mmHg, it is considered a state of critical hypoxia, which can cause irreversible damage within minutes if sustained ^36, 37^. Since the implanted cell mass lacks direct blood vessel perfusion for at least several days, cells located away from the periphery of the graft—and therefore farther from the host’s well-perfused brain tissue—are more prone to hypoxia. A mono-exponential decay model is commonly used to approximate oxygen diffusion in tissue ^38, 39, 40^: *pO*2(*L*) = *pO*2*cap* · *e*^−*L*/*λ*^ (**Supplementary** Fig. 1), where L is the distance from the nearest capillary, *pO*₂*cap* is the initial oxygen partial pressure near the capillary (typically assumed to be 50 mmHg), and λ = 30 µm is the characteristic diffusion length. This model reflects diffusion-limited oxygen delivery, in which oxygen tension drops rapidly with distance from the capillary and approaches critically low values at distances beyond ∼100 µm in metabolically active brain regions ^40, 41^.

To simplify the concept, we can assume that tissue within 100 µm of a capillary is adequately oxygenated. Based on this, the fraction of hypoperfused cells (HypoR) in a spherical implant can be approximated as: 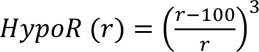 the implanted cell mass increases, a greater proportion of cells are hypoperfused or hypoxic (**Fig. 3H**). No hypoperfusion occurs when r ≤ 100 µm. At r = 100 µm, the volume of the sphere is approximately 4.2 nL, which we define as the threshold volume (Vₜₕ) for the onset of hypoperfusion (**Fig. 3I**). We also calculated the implanted cell mass volume across a range of radii (**Fig. 3I**, upper horizontal axis). For simplified interpretation in the context of cell transplantation experiments, we reformulated the HypoR function using total implanted cell volume instead of radius, 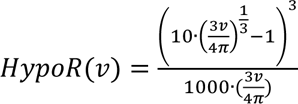 (Equation 4). This produces a skewed bell-shaped curve (**Supplementary** Fig. 2), where the intersection with the x-axis represents Vₜₕ, consistent with the volume threshold shown in **Fig. 3H** (red dashed line).

To validate our mathematical modeling, we conducted experiments to examine how different injection volumes affect cell growth rates. These experiments were performed using a hyaluronic acid (HA) hydrogel matrix, which we previously demonstrated to support neural stem cell graft survival ^42^. C17.2 cells were injected at four distinct volumes into the hydrogel and cultured for 7 days followed by quantification of viable cells using bioluminescent imaging (**Fig. 3J**). We observed that larger injection volumes resulted in more compact, spherical cell masses within the hydrogel, closely resembling in vivo transplantation conditions (**Fig. 3K**). Analysis of the growth rate revealed significantly lower proliferation rates in the 0.5 µL (p = 0.0188), and 1.0 µL (p = 0.0343), groups compared to the 0.1 µL group, while no significant difference was observed between the 0.5 µL and 1.0 µL groups (**Fig. 3L,** p=0.9945). To test our hypoperfusion-based mathematical model, we define the growth rate (G) as a function of the hypoperfused fraction (HypoR): G = Gmax · (1 – HypoR)^3^, where Gmax is the maximal growth rate. As HypoR decreases, G approaches Gmax (**Fig. 3M**). By substituting HypoR with the function derived from cell volume (as shown in **Fig. 3H**), we expressed G as a function of injection volume (v, **Fig. 3M**):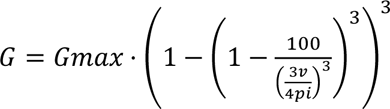 (Equation 5). Using the known *Gmax* value for C17.2 cells (**Supplemental Fig. 3**), we fit the experimental data shown in **Fig. 3L** to the theoretical curve. The resulting fit showed strong agreement, with an R-squared value of 0.78 (**Fig. 3M**). Fitting experimental data at different exponent values also confirms the current model provides the best fitting (**Supplemental Fig. 4**). Thus, under our experimental setting, Equation 5 serves as the hypoperfused fraction prediction function. These results demonstrate that injection volume significantly influences cell proliferation rate. Our mathematical model provides a useful tool for estimating hypoperfused fractions and predicting growth dynamics, offering practical guidance for optimizing cell transplantation protocols.

### OPTRACE enables the development of Pulse-Elevation for Intracortical cell injection

To achieve OPTRACE for in vivo imaging, we intended to leverage intravital TPFM for visualizing donor cells during or after transplantation due to its advantages in spatial and temporal resolution for imaging scattering tissue^43, 44, 45, 46, 47^. As the ideal imaging range of TPMF for mouse brain imaging is around 450 micrometers ^48^, cells need to be implanted in shallow layers (L1-L5) of the mouse cerebral cortex for them to be observable. However, as we learned from the above results, for intracortical transplantation of cells, retention rate will be a huge challenge due to the shallow depths (< 0.5 mm, **Fig. 3E**). To address this issue, we hypothesize that fractionated needle elevation and pulse cell injection with short intervals after each round of needle elevation will improve the retention rate due to the creation of space in needle track after each pulsed injection (**Fig. 4A**).

**Figure. 4.**
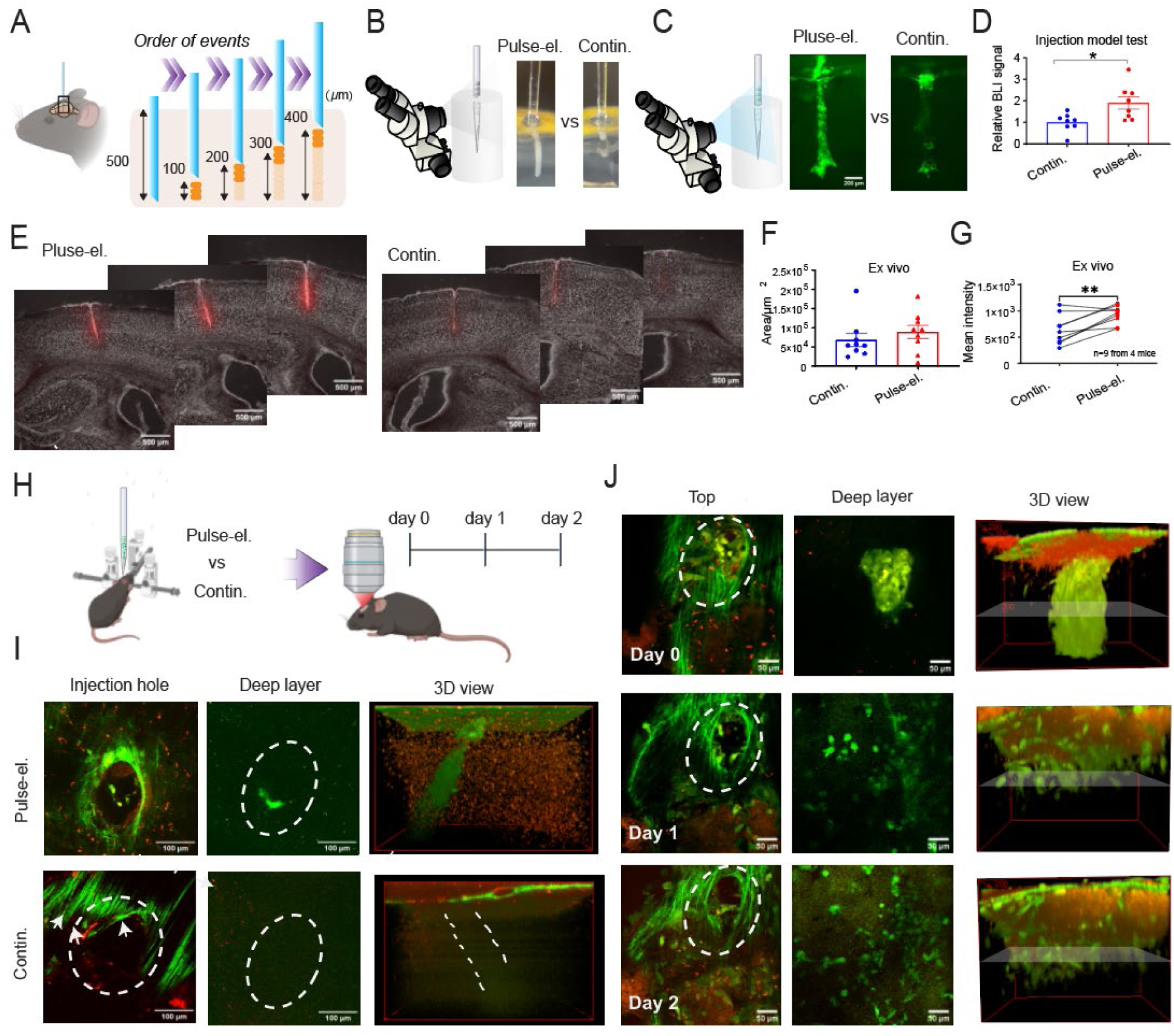
Pulse-elevation model of injection improves cell retention. (**A**) Schematic illustration of the pulse-elevation strategy for transplanting cells into shallow depths of the brain. (**B, C**) In vitro tests under bright field (**B**) or fluorescence (**C**) imaging through a stereoscope. Scale bar, 200 μm. (**D**) Quantitative analysis of bioluminescence signal of C17.2 cells post-injection (*p* = 0.0147, n=8). Differences were analyzed by an unpaired T-test. (**E**) Representative coronal brain slices from Pulse-elevation or Continuous injection groups. (**F**) Quantification of needle track area in continuous and pulse-elevation mode (n=9 from 4 mice). (**G**) Quantification of mean intensity in continuous and pulse-elevation mode (p = 0.006, n=9 from 4 mice). Differences were analyzed by paired T-test. (\**p* < 0.05, \*\**p* < 0.01). (**H**) The experimental design for in vivo TPFM. (**I**) Comparison of the two methods of injection in terms of GFP-positive C17.2 cells residing in needle track imaged right after transplantation under TPFM. Scale bar, 100 µm. (**J**) Daily imaging the transplantation site for three days under TPFM.

We performed *in vitro*, *ex vivo,* and *in vivo* experiments to examine this hypothesis. As we learned that a smaller cell injection volume is preferred to keep the hypoperfused fraction (**Fig. 3H**), we chose 100 nL as the total cell injection volume with a predicted hypoperfusion fraction based on Equation 4 (the predicted value, 0.278). Under the bright field (**Fig. 4B**) or fluorescence imaging (**Fig. 4C**), cells were closely monitored in the continuous or pulse-elevation group, and we observed a significantly high number of cells that were retained in 0.6% agarose gel in the pulse-elevation group (**Fig. 4D**, p=0.0147). Then we confirmed the efficacy of this approach by injecting cells into the mouse brain ex vivo (**Fig. 4E**). Quantitative analysis of the needle track area showed no significant difference between the continuous and pulse-elevation injection modes (**Fig. 4F,** p=0.7344). However, fluorescence signal intensity within the injection tract demonstrated that the mean intensity in the pulse-elevation group was significantly higher than that in the continuous injection group (**Fig. 4G,** p=0.006). This indicates that while both injection modes cause similar levels of tissue damage, the pulse-elevation mode enables greater cell retention within the injection tract. Moreover, the standard deviation of mean intensity in the pulse-elevation group was notably lower (continuous mode=279.9, pulse-elevation mode=170.7, **Fig. 4G**), indicating reduced variability of injected cell number.

We then utilized TPFM to visualize the injection sites immediately after transplantation of C17.2 cells expressing GFP into the mouse cerebral cortex (**Fig. 4H**). A substantial number of cells were retained in the needle track in the pulse-elevation group compared to that in the continuous injection group (**Fig. 4I**). These results indicate that the pulse-elevation mode significantly improves the efficiency of cell delivery, holding promise for optimizing cell-based therapies through minimizing cell loss and enhancing the precision of cell transplantation for therapeutic applications. Next, we extended the observation of implanted cells in the pulse-elevation group through intravital TPFM. On Day 0, the implanted cell mass was visible (**Fig. 4J**, row 1). On Days 1 and 2, a significant shift in cell distribution was observed (**Fig. 4J**, row 2 and 3). While the needle track remained visible, transplanted cells dispersed and migrated away from the injection site. Initially confined to the needle track, the cells were later detected in surrounding brain regions, indicating active migration (**Fig. 4J**, row 3). This pattern suggests that C17.2 cells exhibit intrinsic migratory behavior, likely in response to local microenvironmental cues within the brain tissue, such as signals related to injury, inflammation, or tissue repair.

### A multicolor cell labeling approach for OPTRACE at the Post-Translational Phase

For stem cell therapy, it is highly desirable to shed light into the black box of the post-transplantation behavior of grafted cells from a methodological perspective. Taking advantage of TPFM for high-resolution intravital brain imaging and the availability of genetically encoded reporters, we previously developed an optical cell positioning system ^43^ (oCPS) for intravital long-term single cell tracking of endogenous neuroblasts, based on stochastic mixing of the three colors (Red-Green-Blue, RGB) of fluorescent proteins (FPs, **Fig. 5A**) working similarly to Brainbow ^49^. However, this strategy does not permit functional cell tracking. To solve this problem, we upgraded the RGB marking strategy to ICam (Identity and Calcium) at two aspects (**Fig. 5B**): first, mVenus was replaced by an infrared FP, iRFP682 ^50^ which has a far-red emission easily distinguishable from that of mCherry; second, we include a genetically encoded calcium indicator, GCaMP6s, targeted to the cell nucleus by fusing it to H2B. Thus, the four FPs (mCherry, TagBFP2, iRFP682, H2B-GCaMP6s) used in ICam vectors are spatially and spectrally separated (**Fig. 5B**). We used lentivectors encoding these FPs driven by a constitutive promoter (EF-1alpha) (**Fig. 5C**), which were packaged into lentiviruses (**Supplemental Fig. 5**) for transduction of target cells (primary embryonic neurons or C17.2 cells, **Fig. 5D**) at equal ratio to achieve maximal color variation ^43, 51^, resulting in the production of multicolor-labeled cells with well-scattered color hues (**Fig. 5E**). We also confirm that all four FPs could be detected in transduced cells (embryonic mouse neurons or human iPSC-derived neural progenitors, **Supplementary** Fig. 6A). As for the performance of calcium indicator H2B-GCaMP6s, we induced intracellular calcium rise by mechanical stimulation^52^ (pipetting up and down) and observed a significant rise of fluorescence intensity (p < 0.001, **Fig. 5F and supplementary Fig. 6B**). This confirms the functionality of the calcium indicator in lentivirally transduced cells.

**Fig. 5.**
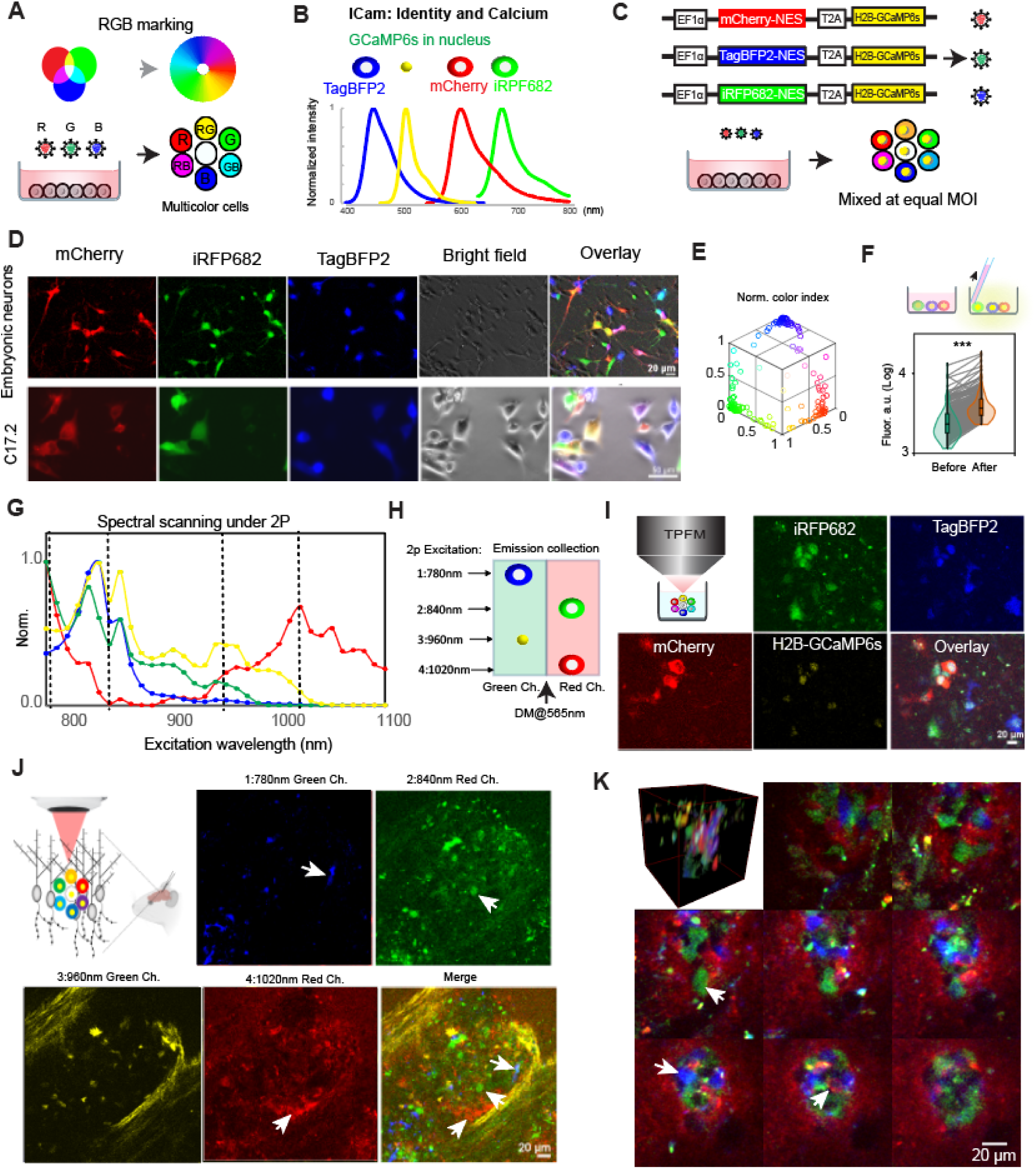
Tracking implanted cells at post-transplantation phase through OPTRACE. (**A**) The principle of RGB marking of cells. (**B**) The component FPs in ICam and their well separated emission spectrum. IRFP682 and H2B-GCaMP6s were false colored as green and yellow, respectively, for better visualization in merged multicolor mode. (**C**) Lentiviral constructs for ICam and the principle of color mixing after cell transduction. (**D**) Primary embryonic neurons from mouse cortex or C17.2 were transduced by ICam lentiviruses at equal MOIs, and the same field-of-view was imaged using different fluorescent filter sets to obtain individual and merged pictures. (**E**) Normalized color index was mapped to 3D-coordinate system for visualization of color distribution. Each dot represents one cell (n = 145 cells). (**F**) Mechanical stimulation was applied by aspirating medium for induction of intracellular calcium rise. Fluorescence intensity of H2B-GCaMP6s expressing cells were compared (p<0.0001, n=263 cells, Krustal-Wallis test). (**G**) 2P excitation spectra for the four FPs used in ICam. (**H**) Election of excitation wavelengths for each FP to unmix their signals. The location of the dot/cells indicates which channel is used for signal collection. DM: Dichroic mirror. (**I**) ICam-labeled hiPSC-NPCs were imaged under TPFM using the setup in (H) for unmixing FP signals. (**J, K**): ICam-labeled cells (hiPSC-NPC, **J**, or C17.2, **K**) were transplanted into mouse cerebral cortex. A representative of xy plane under TPFM showing the ICam labeled cells with separated color channels (arrows, **J**). Montage of the 3D stack of engraft site as a merge of red, green and blue channels showing ICam-labeled C17.2 cells of different colors (arrows) at different depths under intravital TPFM.

As OPTRACE relies on TPFM for cell tracking, we examine if these FPs could be detected and precisely unmixed under TPFM for intravital imaging. HEK293 cells labeled with individual ICam vectors were imaged under TPFM to obtain the two-photon excitation spectra for each FP (**Fig. 5G**). Based on their spectra, we chose for each FP the 2P wavelength that best separates it from others, namely, 780 nm for TagBFP; 840 nm for iRFP682; 960 nm for H2B-GCaMP6s and 1020 nm for mCherry (**Fig. 5H**). This setting enables precise unmixing of these four FPs in multiplex ICam labeled cells (**Fig. 5I and Supplementary** Fig. 7), confirming that we successfully established the labeling strategy that may allow functional single-cell tracking under TPFM. To obtain in vivo proof of the efficacy of our labeling strategy, we utilized the two-photon wavelength setting depicted above to image ICam labeled cells (hiPSC-NPC, **Fig. 5J** or C17.2, **Fig. 5K**) implanted into the mouse cerebral cortex using the pulse-elevation approach. Numerous multicolor donor cells were observed at transplantation sites in either case (arrows). To further evaluate the application of ICam labeled cells for therapeutic use, we transplanted ICam-labeled NSCs (C17.2) into the peri-infarct region (within 0.5 mm to 1 mm from the ischemic core, **Supplementary** Fig. 8A and B) one week after the induction of photothrombosis in the brain ^6, 53^. Two days later, we observed long distance migration of implanted cells from transplantation sites toward the ischemic region (arrows in **Supplementary** Fig. 8C). High magnification observation of the graft reveals the presence of multicolor labeled NSCs with the three morphological color channels (mCherry, iRFP682 and TagBFP2) as well as the nuclear GCaMP6s signal (**Supplementary** Fig. 8D). This result confirms the successful stable multicolor labeling of stem cells with ICam vectors that permits functional single-cell tracking of therapeutic cells.

### OPTRACE at the Post-Translational Phase Reveals Active Interactions Between the Graft and Host Cells

To investigate interactions between implanted cells and the host brain environment, we generated double-transgenic mice by crossing CX3CR1-GFP mice (labeling microglia with GFP) with the GP8.31 line (expressing the red-shifted calcium indicator jReCO1a in neurons) (**Fig. 6A**). The resulting mice exhibited GFP-labeled microglia and jReCO1a-expressing neurons, as confirmed by histology (**Fig. 6B**) and functional calcium imaging (**Supplementary** Fig. 9). We then performed longitudinal imaging from Day 0 to Day 5 after implantation of mouse primary embryonic neurons in the cerebral cortex of this double transgenic mouse line CX3CR1: GP8.31 (**Fig. 6C**). Host neurons (red) surrounding the graft site exhibited a sharp increase in fluorescence intensity shortly after transplantation, which gradually diminished over time (**Fig. 6D**). This pattern was evident in longitudinally tracked neurons located near the graft site, defined by residing within the region of 120 µm radius centered on the implantation site, compared to those positioned beyong the circle (**Fig. 6E**, upper and lower rows, respectively). Quantitative analysis (**Fig. 6F**) showed that calcium signals in the peri-graft region peaked on Day 2 and declined rapidly, returning to near-baseline levels by Day 4. In contrast, neurons distal to the graft consistently displayed low calcium activity throughout the observation period, indicating that the elevated calcium signaling was spatially confined to the immediate graft vicinity. There are significant differences between these two groups on day 0 (p = 0.002), day 1 (p < 0.0001), day 2 (p < 0.0001), and day 3 (p = 0.0105).

**Figure. 6.**
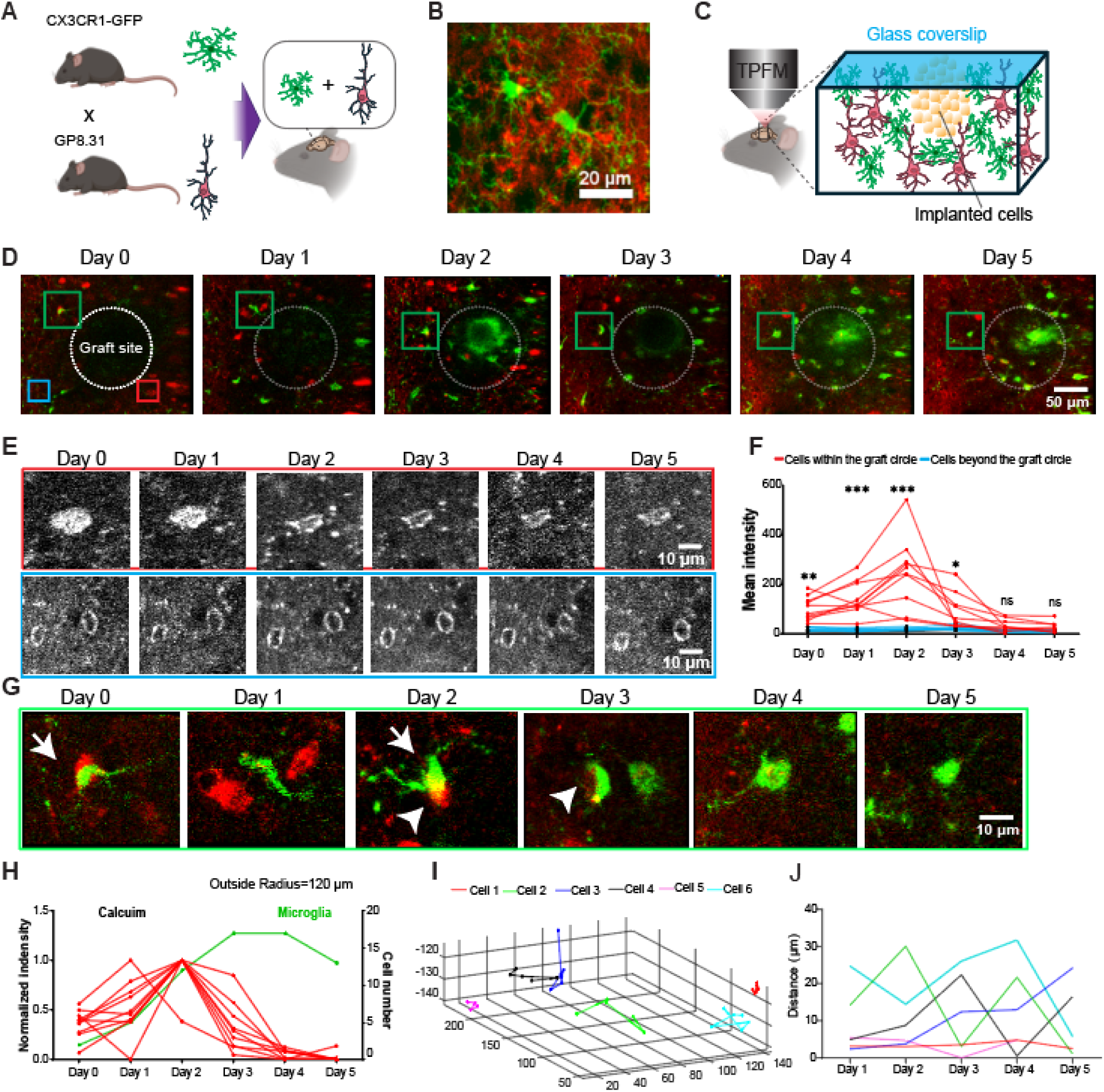
Temporal dynamics of calcium activity and microglial response following cell transplantaion. (**A**) Schematic illustration of the generation of the double transgenic mouse line. (**B**) Example morphology of neurons and microglia in the double transgenic mouse brain. (**C**) Our experimental design: imaging graft-host interactions under TPFM. (**D**) Same area of the brain on different days post-transplantation. Ddashed white circle shows the implantation site with a radius of 120 µm Red tangles indicate regions adjacent to the graft site, whereas blue tangles represent distal regions. Scale bar, 50 µm. (**E**) Magnified views of the same cells shown in (**D**) over time, showing calcium level in cells adjacent to (red box) and distal from (blue box) the graft site over 5 days. Scale bar, 10 µm. (**F**) Quantification of calcium signal intensity in cells located within (red, n = 10 cells) and beyond (blue, n = 8 cells). day 0, p = 0.002, day 1 and day 2, p < 0.0001; day 3, p = 0.0105, day 4, p = 0.436, and day 5, p = 0.5425). Differences were analyzed by two-way ANOVA followed by Tukey’s test. (* p < 0.05, **p < 0.01, ****p < 0.0001). (**G**) An example microglia tracked over 5 days (highlighed in the green tangle shown in **D**). Scale bar, 10 µm. (**H**) Dynamic change of microglia cell number (green) and its relationship to normalized calcium signal (red) within a 120 µm radius surrounding the graft over a 5-day period. (**I**) 3D spatial trajectories of six representative cells within the graft site. Each colored line represents the positional change of an individual cell over time, plotted in three-dimensional space (x, y, z). Cell 2 is the cell shown in (**G**). (**J**) Migration distance of individual microglial cells over a 5-day period. Each colored line represents the tracked movement of a single microglial cell relative to its original position.

In parallel, microglia (green), the brain’s first-line responders to injury, were recruited to the graft area, as indicated by a progressive increase in their numbers within a circular region of 120 µm radius centered on the implantation site (**Fig. 6D** and **6H**). Microglial accumulation peaked around Days 3–4 and declined thereafter (**Fig. 6H**, green line). Interestingly, this response appeared to lag slightly behind the neuronal calcium surge (**Fig. 6H**, red line), suggesting that early neuronal activity may contribute to initiating microglial recruitment. High-resolution immunofluorescence imaging further revealed dynamic interactions between microglia and donor-derived cells (**Fig. 6G**). Microglial processes made direct contact with grafted cells and progressively engulfed cellular debris and damaged neurons (arrows, **Fig. 6G**), consistent with active surveillance and phagocytic engagement in response to the graft. Longitudinal tracking of the same microglia enables us to plot their migration trace in 3D over time (**Fig. 6I**), which displays high cell-to-cell variation in terms of their motility (**Fig. 6J**). This may indicate different functional status requiring further investigation, potentially through molecular profiling techniques, such as scRNA seq.

## Discussion

To overcome long-standing barriers in stem cell transplantation—namely, imprecise delivery and poor visibility into post-engraftment dynamics—we developed OPTRACE: a two-step framework that integrates high-precision, image-guided injection with longitudinal single-cell tracking in vivo. At the time of transplantation, OPTRACE enables real-time visualization of cells through low-cost, translucent glass pipettes under wide-field or TPFM, allowing real-time visualization and optimization of transplantation procedures. This is further enhanced by a novel pulse-elevation injection technique, which improves engraftment in superficial cortical layers. Predictive modeling complements this precision approach by identifying injection parameters that maximize retention and minimize hypoxia stress. After transplantation, OPTRACE leverages multicolor genetic labeling and two-photon imaging to monitor individual grafted cells as well as donor-host interactions over time, revealing key biological processes including donor cell survival and migration, host microglial infiltration, and altered neuronal calcium dynamics at the graft-host interface. By uniting these innovations, OPTRACE offers an accessible and modular platform to not only improve transplantation outcomes but also dissect the complex cellular interactions that underlie graft integration in the living brain.

OPTRACE establishes a new paradigm for intracerebral cell transplantation. Using wide-field or TPFM to visualize fluorescently labeled cells during injection, OPTRACE enables real-time, image-guided delivery of cells with micrometer precision. Our custom apparatus (APP 2.0) – an open-source pipette manipulator upgraded from our previous work ^54^– allows fine control of pipette approach and pressure ejection while continuously imaging the brain (Fig. 1A). This strategy draws on concepts from two-photon–targeted patching (where dyes in the pipette render it visible under 2P)^55^ but extends them to cell transplantation. Notably, all hardware components (imaging scope, micromanipulator, pressure injector) are off-the-shelf or open-source (e.g. APP 2.0), making this approach broadly accessible and adaptable to other laboratories. In short, OPTRACE combines widely available optics and actuators to produce a high-precision transplantation platform, marking a clear advance over prior methods in stem-cell grafting.

Our system also integrates predictive mathematical modeling to optimize transplantation parameters. We derived models relating cell retention to injection depth and hypoperfused tissue volume to injection volume. These models accurately predict outcomes: for example, deeper injections yield higher retention, while larger bolus volumes enlarge the zone of hypoxia. Quantitative simulation thus guides the choice of injection depth and rate that maximizes cell survival and minimize hypoperfusion area. This is important because in prior studies very few injected cells survive long-term – often <5% retention after days^7^– and much cell loss is attributed to physical washout or vascular collapse. In regenerative medicine, such mathematical approaches have been advocated to refine therapy delivery: modeling cell–matrix interactions and perfusion can optimize dosing strategies in silico^56^. Our results exemplify this synergy, showing that predictive models of retention and perfusion can be directly linked to improved engraftment. In practice, OPTRACE users can consult these models to optimize retention rate (Fig. 3F) and improve viability of implanted cells (Fig. 3M). Thus, *the pairing of real-time imaging with a quantitative model* constitutes a powerful innovation for designing transplantation protocols. Future work might extend these models (e.g. to account for tissue compliance or repetitive injections) and integrate them into the OPTRACE software for on-the-fly planning.

A key enabler of OPTRACE is our multicolor and functional cell labeling strategy. By differentially labeling donor cells with combinations of fluorophores and calcium indicators, OPTRACE permits longitudinal functional single-cell resolution tracking. For structural tracking, we previously developed the oCPS strategy^43^ which employ stochastic multicolor labeling, so that each transplanted cells can be uniquely identified by its color hue. Such spectral barcoding is well established in neuroscience: for example, Brainbow and its variants randomly colorize many cells, enabling reconstruction of individual cell morphologies and fates^57^. To add a functional readout, we upgraded the oCPS by incorporating calcium indicators^58^, creating the Identity and Calcium (ICam) constructs, which utilizes spectral and spatial multiplexing for precise identification of each FP. ICam also gives rich color diversity under 2P excitation. This multiplexed labeling dramatically increases throughput compared to monochromatic markers. Two-photon calcium imaging then yields repeated measurements of each cell’s activity. In this way, OPTRACE provides longitudinal structural and functional data: we can follow how specific grafted neurons survive, migrate, and become active (or silent) over time. Such capability contrasts sharply with traditional end-point histology or static imaging. Two-photon calcium imaging is a powerful approach to measure neuronal activity at high spatial resolution in neuroscience field^45, 46^ and our use of GCaMP within transplanted cells extends this to the grafted population. In short, OPTRACE’s palette of labels (multicolor fluorophores plus functional indicator) combined with 2P microscopy offers an unprecedented window into the post-engraftment life of single cells.

Using this platform, we uncovered dynamic graft–host interactions. For example, we observed marked changes in host neuron calcium dynamics after transplantation. Immediately adjacent to the graft site, many host neurons showed altered calcium transient frequency and amplitude compared to remote cortex, suggesting that the injection and presence of new cells modulate local network activity. These perturbations in neuronal calcium signaling may reflect tissue injury, synaptic remodeling, or inflammation, and they bear on functional integration. Importantly, we also tracked the immune response: host microglia rapidly invaded the graft. In vivo two-photon imaging revealed host-derived microglial migrate into the transplant within days, consistent with literature^59^. Through OPTRACE, we were able to track the migration of microglia, engulfing neurons with high calcium level and cleaning the debris (Fig. 6). These observations suggest that microglia are actively sensing and responding to the new cells and any perturbations they induce. Furthermore, we found that neurons close to transplantation site displayed elevated calcium activity. Several in vivo two-photon calcium imaging studies consistently demonstrate that neurons near sites of brain injury or implantation exhibit elevated calcium activity, while neurons farther away remain relatively unaffected. For example, Eles et al. (2018) showed that cortical neurons within ∼150 μm of an implanted electrode displayed acute calcium overload, unlike distant neurons ^60^. Similarly, Bibineyshvili et al. (2024) reported a localized calcium surge near a focal cortical injury with suppressed activity in remote regions^61^. These findings align with our observation that neuronal calcium activity is heightened in the vicinity of the transplantation site, supporting the notion of spatially restricted excitability following cortical disruption. Overall, OPTRACE reveals a rich picture of graft integration: donor neurons gradually mature and form functional connections, while host microglia dynamically infiltrate, surveil, and possibly prune graft tissue. Such *live-cell evidence* of neuronal and glial interplay could only be obtained with in vivo tracking; it provides mechanistic insight into why some grafts succeed or fail.

Compared to existing stem-cell transplantation monitoring paradigms, OPTRACE offers major advantages in resolution, precision, and insight. Conventional approaches have primarily relied on post-hoc histology or low-resolution imaging. For example, many studies including ours have used bioluminescent reporters (e.g. luciferase) to monitor graft survival noninvasively^42, 62, 63^. While such methods can report whether transplanted cells are active on a whole-animal level, they lack cellular detail. Likewise, magnetic resonance or PET imaging can track cell location macroscopically but cannot resolve individual cells or acute dynamics. TPFM has been used for observation after transplantation ^59, 64, 65, 66, 67, 68^, but the uniform cell labeling makes it hard to track highly dynamic cells over time. In contrast, OPTRACE provides high accuracy through multicolor cell labeling and couples imaging to the delivery itself. By seeing the pipette in real time, we dramatically reduce targeting errors: empirical recovery rates of transplanted cells are higher (e.g. 10–20% of cells remain versus <5% normally ^7^. Moreover, our spatial resolution is unparalleled: OPTRACE pinpoints cells to within tens of microns, and can track individual cells over time, something not possible with bulk imaging or single-color labels. Temporally, OPTRACE is effectively continuous (images acquired many times per second during injection and repeatedly afterwards), whereas modalities like bioluminescence integrate signals over minutes to hours. In summary, OPTRACE merges the cell-level precision of microscopy with transplantation, giving both space- and time-resolved control and readout.

In conclusion, OPTRACE represents a step-change in how we perform and study neural cell transplantation. By integrating accessible imaging hardware and modeling, it achieves levels of precision and quantitative control previously unattainable. Our findings of graft-host neuronal and immune dynamics open new avenues for optimizing cell therapies. Future work might extend OPTRACE to other cell types (e.g. oligodendrocyte progenitors or CAR-T cells), explore optogenetic control in transplanted networks, or integrate magnetic resonance or ultrasound for even deeper guidance. Importantly, the open design (APP 2.0) invites the community to adapt OPTRACE widely, accelerating progress. Ultimately, by pairing live optical readout with injection, OPTRACE not only improves engraftment outcomes, but also provides a rich experimental platform – a “living microscope” – for studying the biology of transplanted cells in situ.

## Materials and Methods

### Glass pipette preparation

Glass pipettes were fabricated from borosilicate capillaries (Drummond Scientific Wiretrol II / Plungers - WIRETROL II, Cat#1-5 UL - 5-000-2005) using an affordable pipette puller. The pipette tips were subsequently beveled using a custom-built rotary grinder adapted from an external hard drive motor ^27^. Pipettes were ultrasonically cleaned in 1 M NaOH (Sigma Aldrich, Cat#S5881) for 15 minutes and then rinsed twice in double-distilled water, each for 15 minutes. Residual water inside the pipettes was removed by vacuum aspiration using a fine-tip adapter. The dried pipettes were subsequently filled with mineral oil (Sigma Aldrich, Cat#M8410) using a backfilling technique and connected to a 10 μL Hamilton micro syringe (Hamilton, model 1701N) via a steel adapter. All pipettes were prepared on the day of use.

### Cell cultures

HEK-293 cells were cultured under identical conditions using DMEM supplemented with 10% fetal bovine serum (FBS, Sigma, Cat#F4135), 2 mM L-glutmax (Gibco, Cat#35050-061), and 1% penicillin–streptomycin (P/S; Sigma, P433). Mouse neural progenitor cells stably expressing *LacZ* (C17.2-luc2⁺) ^63^ were maintained in high-glucose Dulbecco’s Modified Eagle Medium (DMEM,Gibco,Cat#11965118) supplemented with 10% FBS, 5% horse serum (Gibco, Cat#26050-070), and 1% P/S at 37 °C in a humidified atmosphere containing 5% CO₂. Primary cortical neurons were isolated from embryonic day 17.5 C57/BL6 mouse embryos, as previously described ^69^. Briefly, cortices were dissected and digested in 0.25% trypsin-EDTA (ThermoFisher, Cat#25200056) containing 100 U/mL DNase I (Sigma Aldrich, Cat# 10104159001) at 37 °C for 15 minutes, with gentle agitation every 5 minutes. Following enzymatic digestion, the tissue was gently triturated several times to dissociate aggregated tissue. After allowing the debris to settle for 2 minutes, the supernatant containing dissociated cells was transferred to a 15 mL centrifuge tube and centrifuged. The resulting cell pellet was resuspended in DMEM supplemented with 10% FBS and plated at a density of 1 × 10⁶ cells per well in 6-well plates containing poly-D-lysine coated (50 μg/mL, ThermoFisher, Cat# A3890401). Cells were initially cultured in DMEM with 10% FBS for 2 hours to facilitate cell attachment. The medium was then replaced with Neurobasal medium (Thermo Scientific, Cat#21103049) supplemented with 2% B27 (Thermo Scientific, Cat#17504044). Neurons were maintained at 37 °C in a humidified incubator with 5% CO₂. Human iPSC-derived neural precursor cells (NPCs, Elixirgen Scientific, Inc. CIRM line CW50065) were cultured following the Quick-Neuron Precursor protocol (Elixirgen Scientific, Inc.). Cells were thawed, centrifuged, and plated onto 6-well plates pre-coated with Poly-L-Ornithine and laminin in NPC Medium (A). Cultures were maintained in NPC Medium under standard conditions (37°C, 5% CO₂) with media changes on days 1, 2, and 4. On day 7, cells were either passaged using a TrypLE or cryopreserved.

### Cell transduction by lentivirus

All lentivectors were cloned and packaged by VectorBuilder and tittered by us. For multicolor lentiviral transduction, target cells were transduced with lentiviruses encoding distinct fluorescent proteins (e.g., iRFP-682, mCherry, and BFP) at MOIs indicated in the results. The viral mix was supplemented with 5 μg/mL polybrene (VectorBuilder) to enhance transduction efficiency. After 24 hours of incubation with the viral supernatant, the medium was replaced with fresh medium. Transduction efficiency was assessed by 72 hours post-infection using flow cytometry (BD LSR II) or florescence microscopy (Leica DMi8). For long-term expression and lineage tracing, cells were sorted based on fluorescence profiles via fluorescence-activated cell sorting (FACS, BD Aria II) expanded and maintained under standard conditions for downstream applications.

### Spectral scanning under two-photon imaging

Hek-293 individually labeled with ICam lentivectors were imaged under two photons at excitation wavelength ranging from 780-1100 nm at step size 10 nm. Emissions were split by a dichroic mirror (565DCXR, Chroma) and detected by PMTs H16201P-40//004, Hamamatsu) into green and red channel detected by PMTs. ROIs were drawn on the same area of images to measure intensity. After background subtraction, intensity was normalized to 0 to 1 based on minimal and maximum values.

### Cell viability test after flowing through glass pipettes

Following trypsinization with TrypLE (Gibco, Cat#12604021), cells were centrifuged at 1000 rpm for 5 min and resuspended at the target density in PBS (phosphate-buffered saline, pH 7.4) (Gibco, Cat#10010-023) supplemented with 1% bovine serum albumin (BSA, Sigma, Cat#A8412) and 0.5 mM EDTA (Ethylenediaminetetraacetic acid; Quality Biological, Cat#351-027-721). Cells were pipetted ten times per injection, with pipetted cells serving as the control group. For glass pipette diameter selection experiments, 2 μL of cell suspension (2.5 × 10⁷ cells/mL) was injected into 18 μL of 0.4% Trypan Blue Solution (Thermo Fisher, Cat#15250061), gently mixed, and immediately analyzed using the Cellometer Auto 2000 Cell Viability Counter (Nexcelom Bioscience).

### Preparation of brain simulation scaffold for injection depth and volume testing

For injection depth testing, cell suspensions (2.5 × 10⁷ cells/mL) were injected at defined depths of 0.5 mm, 1.5 mm, and 3.0 mm into 0.6% low melting point agarose (Promega, Cat#v2111) using a glass pipette with a 100 µm tip diameter. A total volume of 1 µL was injected at a constant rate of 0.5 µL/min, waiting for 2 mins before withdrawing. Following injection, cell viability within the agarose constructs was immediately assessed using bioluminescence imaging (BLI). For injection volume test, HyStem-HP hydrogel (Sigma Aldrich, Cat#HYSHP020) was prepared according to the manufacturer’s instructions for use in in vitro injection volume testing. To form the hydrogel, equal volumes of HyStem-HP and Gelin-S were mixed to obtain a 1:1 polymer blend. A total of 2.0 mL of this blend was then combined with 0.5 mL of 1× Extralink2 stock solution (resulting in a 4:1 polymer-to-crosslinker ratio, v/v), and the mixture was homogenized by gentle pipetting. Subsequently, 70 µL of the final pre-gel solution was loaded into a custom gel holder fabricated from a trimmed 1 mL pipette tip (top inner diameter: 2 mm; bottom inner diameter: 5 mm; height: 9 mm). The filled holders were incubated at room temperature for 20–30 minutes to allow complete gelation. A series of defined volumes (0.1 µL, 0.25 µL, 0.5 µL, and 1 µL) of cell suspension were injected into the hydrogel using a glass pipette with a 100 µm tip diameter. Injections were performed at a constant rate of 0.25 µL/min to a depth of 2.3 mm. Following injection, cells were transferred to 96-well plate in incubator and cell viability within the hydrogel constructs was evaluated on 0, 3, and 7 days post-injection using BLI. At each time point, D-luciferin (Gold Biotechnology, eLUCK-100; 15 μg/mL in 10 mM PBS, pH 7.4) was added directly to each well containing cells. Luminescent signals were acquired every 10 minutes using the Spark CYTO (Tecan). Peak luminescence values were subsequently used for quantitative analysis of cell survival.

### Pulse-elevation mode injection into agarose gel and quantification

C17.2-luc2⁺ cells (2.5 × 10⁷ cells/mL) were injected into 0.8% low-melting-point agarose. The target depth for the cell transplantation was set at 500 μm. Before injection, glass pipette was extended beyond the target depth to 1000 µm and then retracted back to a target depth of 500 µm. For the pulse-elevation mode, a five-step infusion protocol was employed to deliver a total volume of 100 nL at a constant rate of 0.1 μL/min. A total of 20 nL was injected at sequential depths of 0.5, 0.4, 0.3, 0.2, and 0.1 mm, with a 20-second pause between steps. During each injection, the glass pipette was gradually retracted, and after the final step at 0.1 mm, it was held in place for 5 minutes to promote diffusion and reduce backflow. In the continuous mode, the same cell suspension, depth, injection rate (0.1 μL/min), and total volume (100 nL) were used, but the entire volume was delivered in a single uninterrupted step. Retained cells were quantified using bioluminescence imaging (BLI).

### Comparison of injection methods into dissected mouse brain and analysis

HEK-293 cells (3.0 × 10⁷ cells/mL) were labeled with 25 μM Vybrant™ DiI cell-labeling solution (Invitrogen™, Cat#V22889) following the manufacturer’s protocol. Briefly, 5 mL of 25 μM DiI solution was added to the cell pellet after enzymatic digestion and centrifugation. The cell suspension was gently mixed by pipetting and incubated at 37 °C for 20 minutes. After incubation, cells were centrifuged and washed three times with sterile PBS to remove excess dye. The labeled cells were then resuspended to a final concentration of 2.5 × 10⁷ cells/mL. A total volume of 100 nL of DiI-labeled HEK-293 cell suspension was injected into predefined brain regions (ML: ±1.5 mm; AP: ±1.2 mm and ±2.5 mm; DV: –0.5 mm) at a constant rate of 0.1 μL/min. Injections were performed using two modes: pulse-elevation mode for the right hemisphere and continuous mode for the left hemisphere. Immediately following the injection, brains were fixed in 4% paraformaldehyde and cryoprotected in 30% sucrose in PBS. Coronal brain sections (30 μm thickness) were prepared using a cryostat (CryoStar NX50, Epredia) and imaged using a fluorescence microscope (Leica DMi8) to evaluate the distribution of injected cells.

For ex vivo analysis, 25 μm-thick brain slices containing DiI-labeled transplanted cells were imaged using a Leica fluorescence microscope. All images were acquired under identical settings, including exposure time and gain, to ensure consistency across samples. The images were imported into ImageJ (Fiji), and brightness and contrast were adjusted uniformly using the “Brightness/Contrast” tool. To quantify the injection site, the DiI-labeled injection track was manually outlined using the “Freehand Selection” tool, and the selected region of interest (ROI) was added to the ROI Manager. The area and mean fluorescence intensity within each injection track were measured and exported for further analysis.

### Mice

All experimental protocols at the University of Maryland, Baltimore, were conducted according to the National Institutes of Health guidelines for animal research and approved by the Institutional Animal Care and Use Committee at the University of Maryland, Baltimore. Mice were group housed with littermates until craniotomy surgery, after which they were singly housed. Mice were maintained on a 12–12-h (6 a.m.–6 p.m.) light–dark cycle.

### Cell transplantation and cranial window installation

Male C57BL/6J mice or double transgenic mice breed from CX3CR1-GFP (JAX mice strain # 005582) and GP8.31(JAX mice strain #030526) were anesthetized with isoflurane (5% for induction and 1.5% for maintenance) and positioned in a stereotaxic frame (RWD Stereotaxic Instruments). A circular craniotomy (3 mm diameter) was performed over the primary visual cortex (V1), centered 2.5 mm lateral and 0.5 mm anterior to the Lambda suture. A 100 nL volume of the cell suspension (2.5×10⁷ cells/ml) was injected into the cortex at coordinates: AP = −2.5 mm, ML = 2.5 mm, DV = 0.5 mm using a 100 μm ID glass pipette at a rate of 0.1 μL/min. After injection, a 3-mm diameter circular coverslip adhered to a 3.5-mm diameter donut-shaped coverslip (Assistent Deckgläser Microscope Cover Glasses) was affixed over the craniotomy using black dental cement (Contemporary Ortho-Jet). A custom titanium headpost was then secured to the skull with dental cement. Mice were allowed to recover on a heating pad before being returned to their home cages.

### Two-photon imaging and analysis

Mice were kept on a warm blanket (37 °C) and anesthetized using 0.5% isoflurane and Imaging was performed with a custom-built two-photon microscope with a resonant scanner. Each experimental session lasted 45 min to 2 h. Multiple sections (imaging planes) may be imaged within the same mouse. Fluorophores were excited by a different wavelength with a femtosecond laser system (Chameleon Discovery, Coherent) that was focused by an Olympus ×25, 1.05-NA objective. Emitted fluorescence photons reflected off a dichroic longpass beam splitter (FF705-Di01–25×36, Semrock) and were split with a dichroic mirror (565DCXR, Chroma) and detected by PMTs H16201P-40//004, Hamamatsu) after filtering with a 510/84-nm filter (84-097, Edmund) for the green channel and two 750SP filters (64-332, Edmund) for the red channel. Images were acquired using ScanImage^65^ (Vidrio Technologies). Three-dimensional z-stack images were acquired at a step size of 2 µm along the z-axis. After imaging, animals were returned to their home cages for recovery. For longitudinal imaging at subsequent time points, the same imaging parameters and field of view were used for data acquisition from the same animal. In mice transplanted with multicolor-labeled cells, wavelength optimization was first performed to determine the most effective excitation conditions for each fluorophore. Based on the results, specific wavelengths were used for each channel: 780 nm for BFP, 842 nm for iRFP-682, and 1020 nm for mCherry and GcaMP6s. The laser power ranges from 4 to 16.8 mW.

For real-time imaging cell injection process into the dissected mouse brain, C17.2 cells labeled with GFP (2.5 x10^7^/mL) were injected through glass pipette (ID=110 μm) under two-photon imaging with excitation at 960 nm. The dissected mouse brain was immersed in cold artificial cerebrospinal fluid CSF (aCSF) on ice. ACSF was prepared by dissolving 124 mM NaCl, 3 mM KCl, 1.25 mM NaH₂PO₄·H₂O, 1 mM MgSO₄·7H₂O, 2 mM CaCl₂·2H₂O, 26 mM NaHCO₃, and 10 mM D-glucose in distilled water. After adjusting the final volume to 1 L with distilled water, the aCSF was used immediately or stored at 4 °C for short-term use.)

Imaging data were processed with custom programs written in MATLAB (Mathworks) and Fiji^70^. Raw two-photon imaging data were imported into ImageJ for the generation of composite images. Three-dimensional image reconstructions were obtained using the “3D Viewer”. Fluorescence intensity of calcium signals in local neurons was quantified by drawing ROIs over the soma of neurons and the background was subtracted. For microglial tracking, first, we identified the same cell across time points using consistent local anatomical landmarks and stable reference cells, including non-activated local neurons and stationary neighboring cells. Then we mapped their location into one coordinate system. The x, y, and z coordinates of the same microglial cell were recorded across time within the same field of view. Spatial migration and migration distance of microglia were then analyzed using custom-written scripts in MATLAB.

### Statistical analysis

Data analysis was performed using a combination of standard functions and custom scripts in MATLAB, Prism 10 (GraphPad Software Inc. US) or PAST^71^. The data were tested for normal distribution using the Shapiro–Wilk test. Parametric tests were used for normally distributed data, and non-parametric tests were applied to all other data. Bar graphs and mean ± SEM were used to describe the data with normal distribution, while boxplots and median ± IQR were used to describe the non-normally distributed data. Boxplots represent median and 25th - 75th percentiles, and their whiskers are shown in Tukey style (plus or minus 1.5 times IQR). For comparisons among multiple groups under a single factor, if the data were normally distributed, one-way ANOVA followed by Turkey’s test was performed. If data were not normal distributed, a nonparametric test (Wilcoxon signed rank test) was used to examine paired data (Figure 4F). The statistical significance was defined as ∗p < 0.05, ∗∗p < 0.01, ∗∗∗p < 0.001, < 0.0001, respectively. Medians, IQR, means and SEM are reported throughout the text.

## Acknowledgements

Y.L. acknowledges funding support from the National Institutes of Health (R21AG077631; R03NS123733; R03NS128459; R21AG074978) and Maryland Stem Cell Research Fund (2024-MSCRFD-6363 and 2022-MSCRFL-5893). P.W. acknowledges funding support from Maryland Stem Cell Research Fund (2022-MSCRFD-5886) and NIH (R01DA056739). J. W. acknowledges funding support from Maryland Stem Cell Research Fund for the postdoc fellowship (2024-MSCRFF-6328). We acknowledge the Confocal Microscopy Core Facility, part of the Center for Innovative Biomedical Resources (CIBR) at the University of Maryland School of Medicine, for access to imaging instrumentation and expertise. The authors also thank Flow Cytometry Shared Service of the University of Maryland Marlene and Stewart Greenebaum Comprehensive Cancer Center. Illustrative elements in this manuscript were created using academic resources from BioRender (www.biorender.com), BioArt (https://bioart.niaid.nih.gov/), and Bioicons (www.bioicons.com). We gratefully acknowledge the component library developed by Alexander Franzen (https://www.gwoptics.org/ComponentLibrary/), which was used for the illustration in Figure 2.

## Author contributions

Y.L. designed the research. W.J. performed cell culture, ex vivo and in vivo imaging experiments. C.R., M. W. and D. G. designed and tested the upgraded APP. G. Q, C. C, M. J and P.W. contributed to design and imaging of animal experiments. F.X. contributed to flow cytometry. Y. L. and P.W. supervised research. Y.L. and W.J. wrote the manuscript.

## Materials & Correspondence

Design files: design files are available here: https://github.com/liangy10/APP2. All data are available from the Lead Contact, Yajie Liang, upon request.

