## Supplementary Figures for "OPTRACE: Optical Imaging–Guided Transplantation and Tracking of Cells in the Mouse Brain"

**Supplementary Figure 1-9**

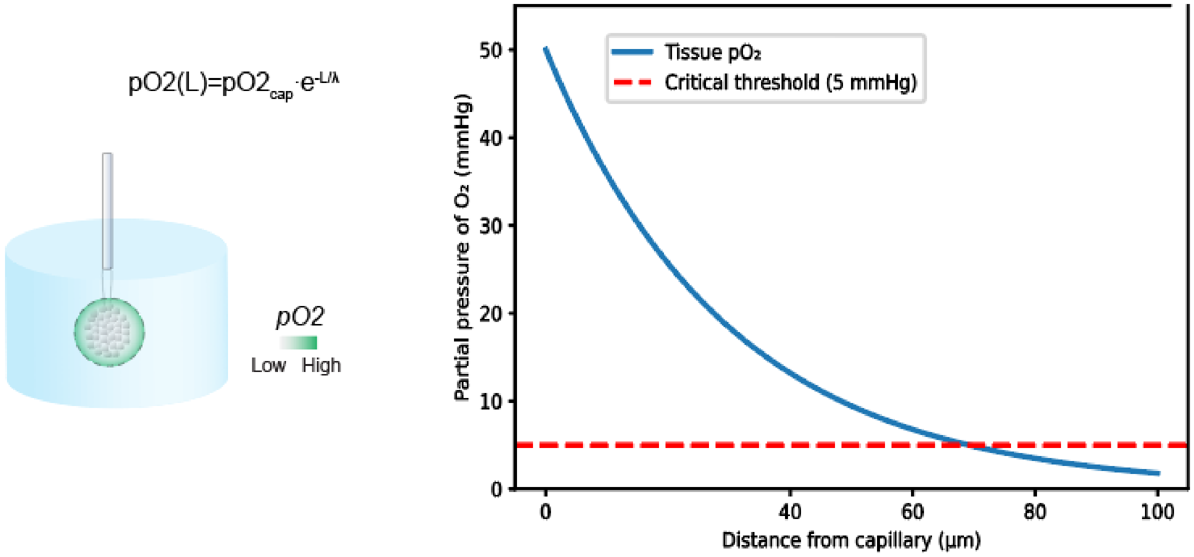


**Supplementary Figure 1. The mono-exponential decay model for evaluating oxygen diffusion in tissue.** L is the distance from the nearest capillary, $pO_2cap$ is the initial oxygen partial pressure near the capillary (typically assumed to be 50 mmHg), and λ = 30 μm is the characteristic diffusion length. This model reflects diffusion-limited oxygen delivery, in which oxygen tension drops rapidly with distance from the capillary and approaches critically low values at distances beyond ~100 μm.

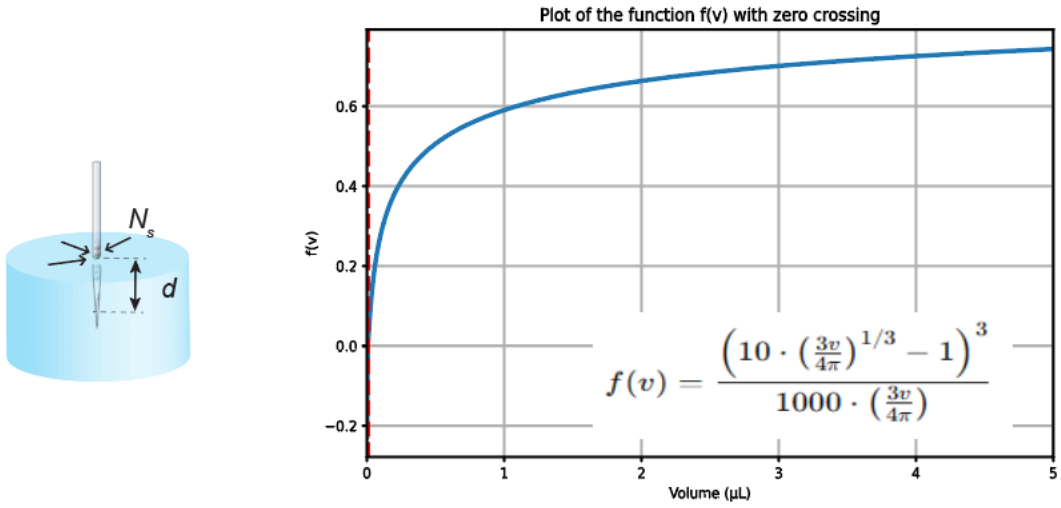


**Supplementary Figure 2. HypoR as a function of implanted cell volume.** The intersection with the x-axis represents $V_{th}$.

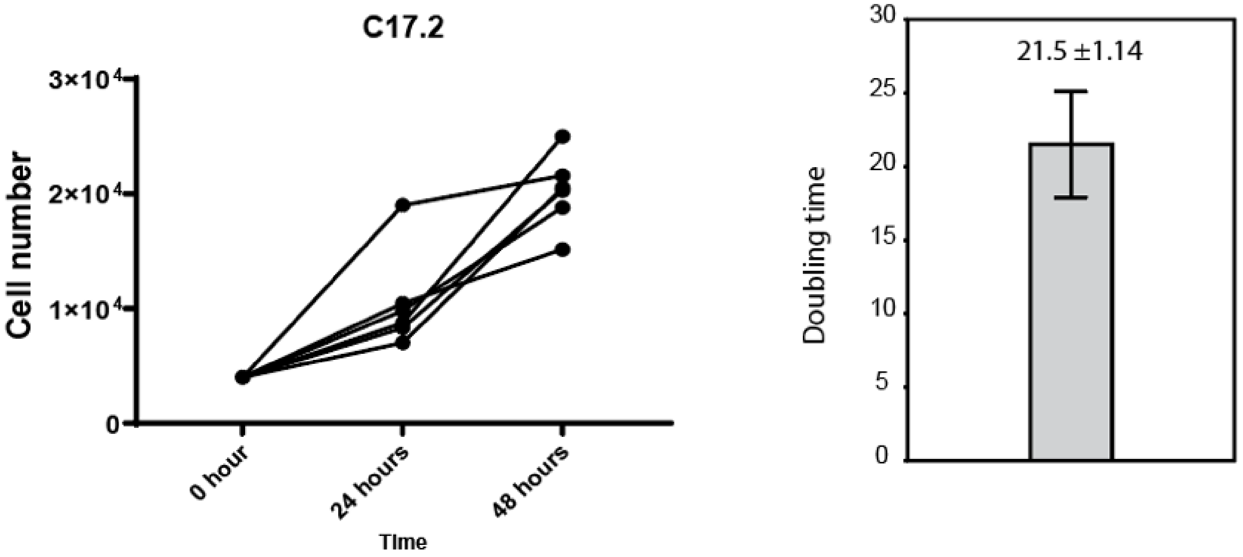


**Supplementary Figure 3. Determination of Gmax value for C17.2.** Cells were cultured in tissue culture plate and cell number was quantified by cck8 at different timepoints. Doubling time was calculated by Time x Log(2)/log(Nt/N0), in which Nt as cell number at Time t and N0 as starting cell number.

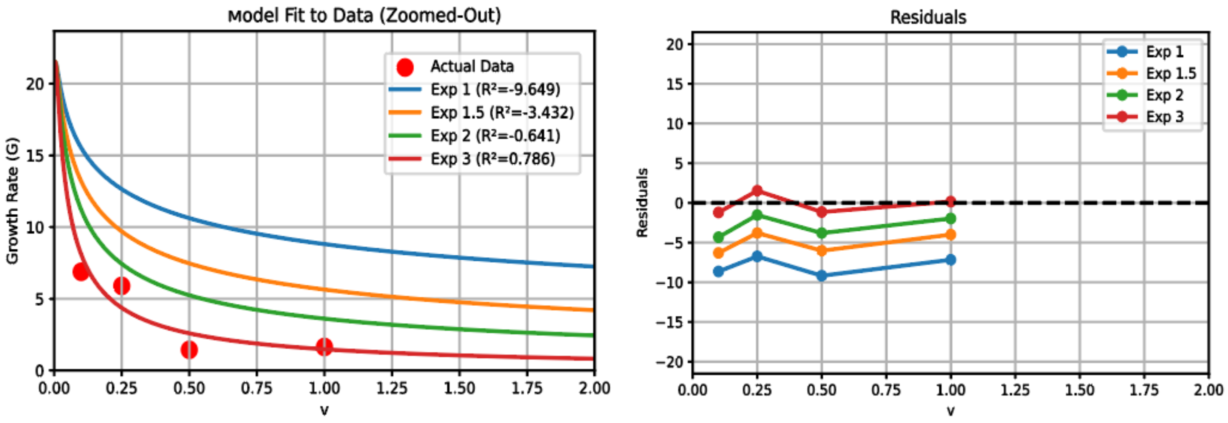


**Supplementary Figure 4. Fitting experimental data to formula with different exponential.** Exponential at 3 gives best fitting evidenced by largest R square value (left panel) and least residual (right panel)

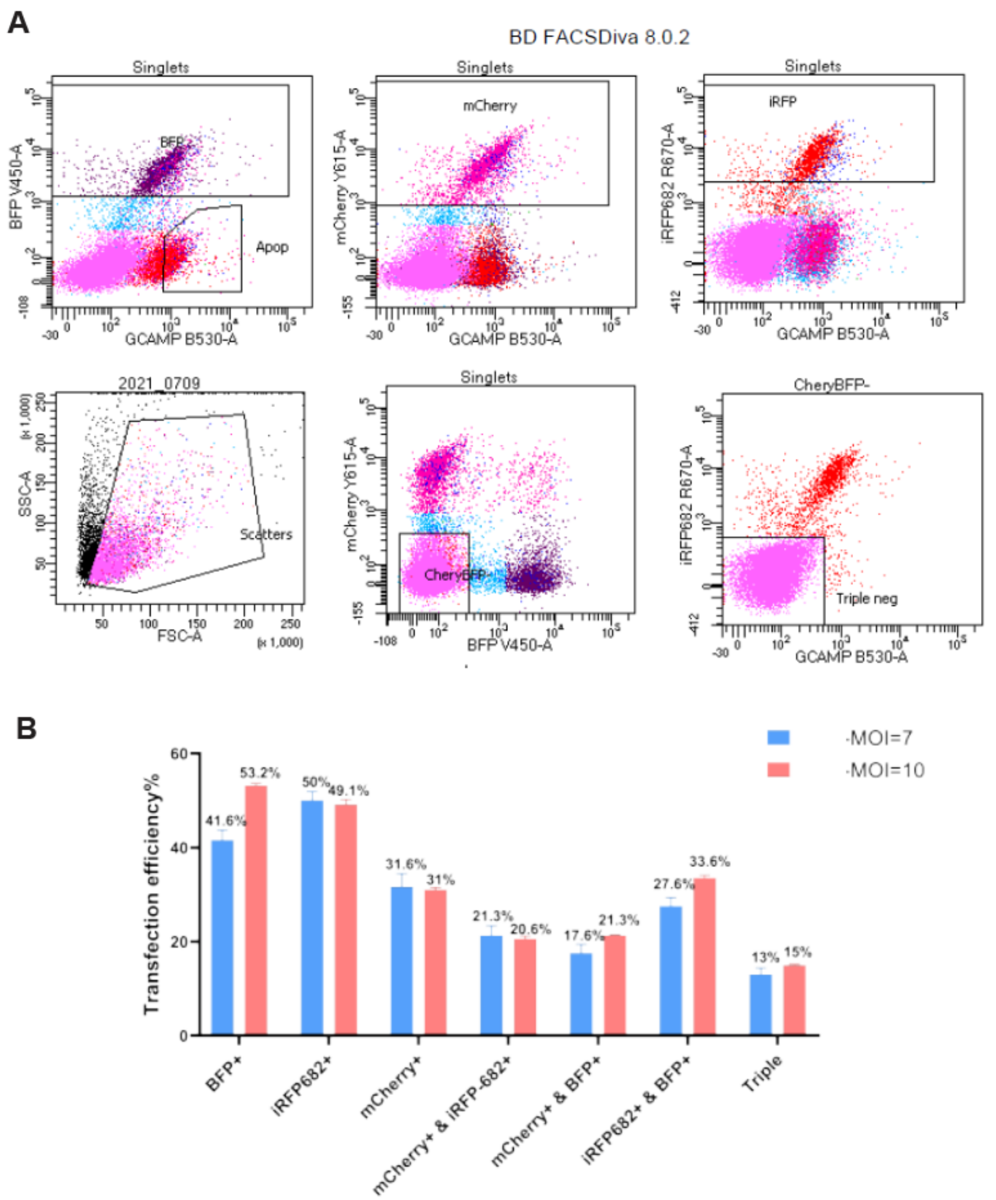


**Supplementary Figure 5. Characterization of ICam lentivectors for mixed transduction of HEK293 cells. A**. All four FPs in ICam vectors were readily detected through flow cytometry. **B**. Quantification of transfection efficiency for different populations of cells labeled with single, double or triple colors in transduced HEK293 cells.

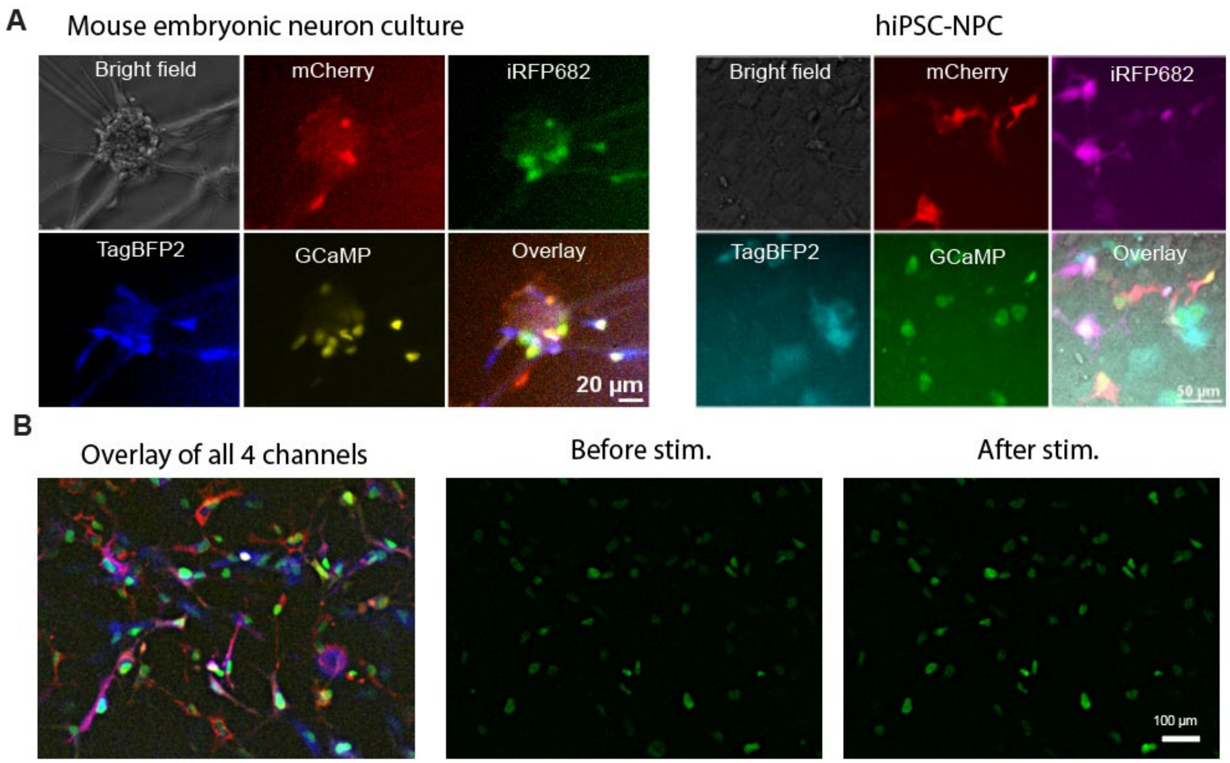


**Supplementary Figure 6. Confirmation of FPs and calcium indicator functionality. A**. All four FPs in ICam vectors were readily detected in transduced cells (embryonic mouse neurons or human iPSC-derived neural progenitors, NPC). **B**. In ICam transduced iPSC-NPC, mechanical stimulation induced the rise of green fluorescence (Right two panels).

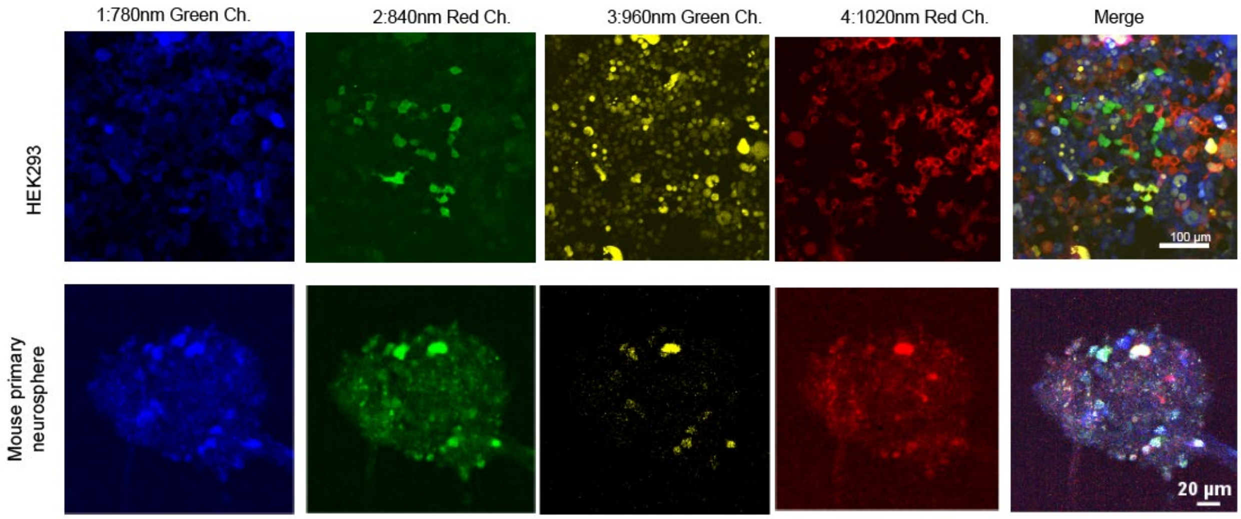


**Supplementary Figure 7. 2P acquisition settings.** Unmixed FP signals were extracted using our special excitation wavelength set in multiplex ICam labeled HEK293 or mouse primary neurospheres.

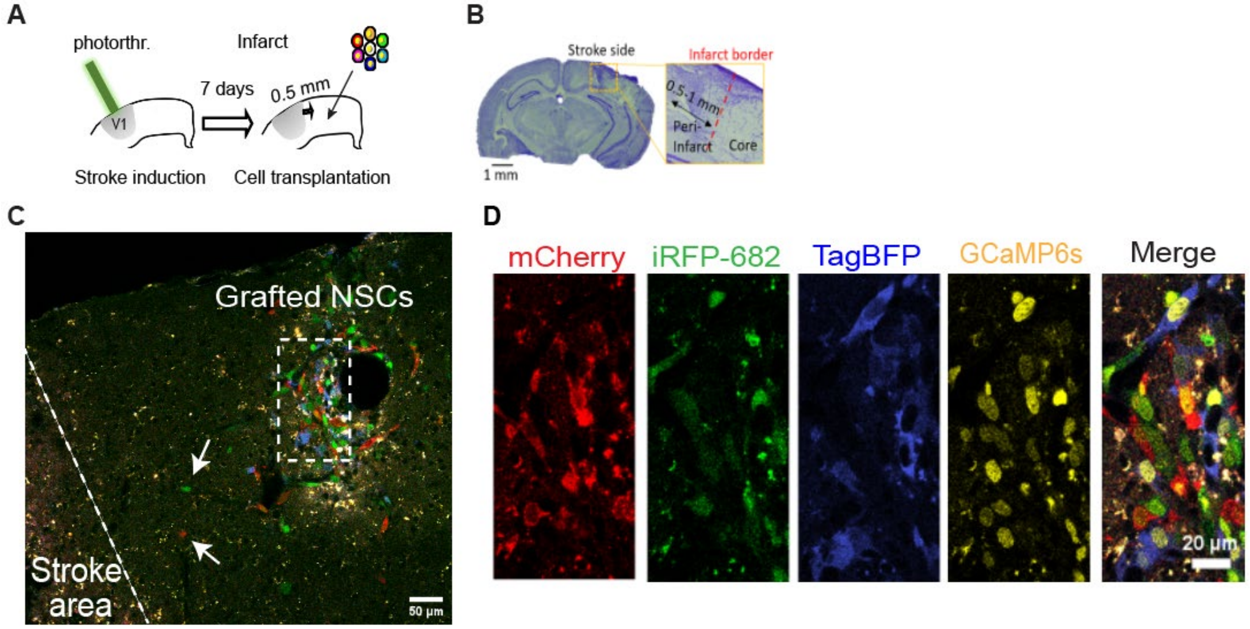


**Supplementary Figure 8. Using ICam labeled cells for therapy of a stroke model.** A, B. Photothrombosis was induced (A) one week before the transplantation of ICam-labeled C17.2 cells into peri-infarct region (B). C. Migration of ICam-labeled NSCs (arrows) toward ischemic region two days after transplantation. D. Zoom in of dashed region in (C) with the breakdown of color channels.

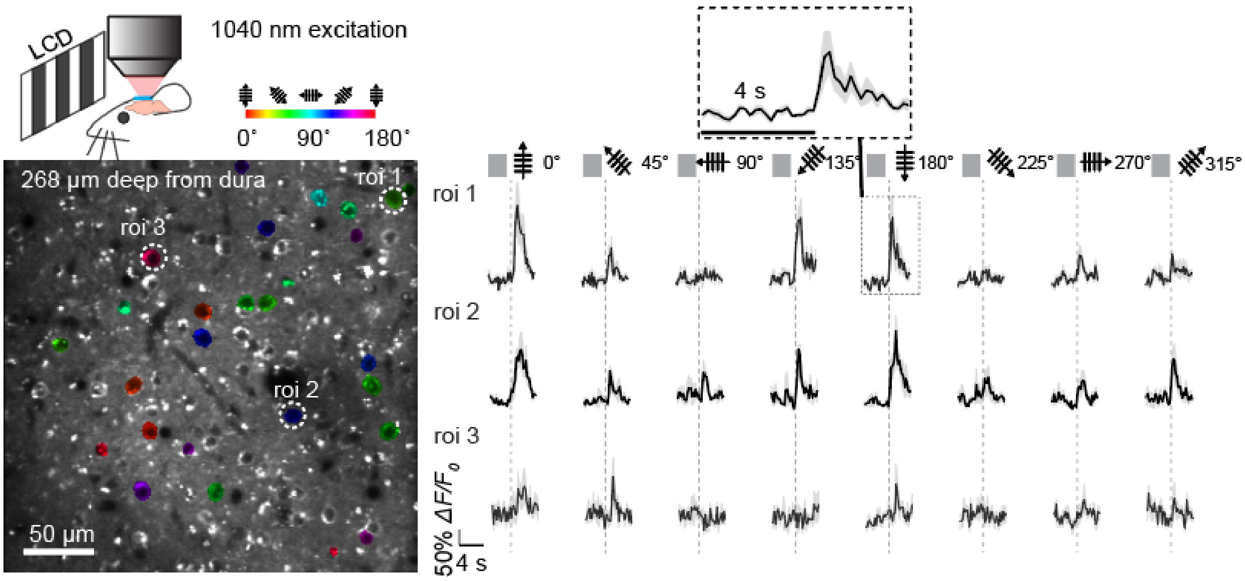


**Supplementary Figure 9. Calcium responses from neurons expressing jRGECO1a.** We imaged Layer 2/3 cortical neurons in the mouse primary visual cortex with 1040 nm 2-photon laser while the animal was exposed to drifting grating visual stimulation 3 weeks after cranial window installation. Cortical neurons in V1 show clear orientation selectivity (left panel) and there are big-magnitude responses from these neurons that could be as high as 150 ΔF/F% above base line (right panel).
